# DHS-Crystallize: Deep-Hybrid-Sequence based method for predicting protein Crystallization

**DOI:** 10.1101/2020.11.13.381301

**Authors:** Azadeh Alavi, David B. Ascher

## Abstract

The key method for determining the structure of a protein to date is X-ray crystallography, which is a very expensive technique that suffers from high attrition rate. On the contrary, a sequence-based predictor that is capable of accurately determining protein crystallization property, would not only overcome such limitations, but also would reduce the trial-and-error settings required to perform crystallization. In this work, to predict protein crystallizability, we have developed a novel sequence-based hybrid method that employs two separate, yet fully automated, concepts for extracting features from protein sequences. Specifically, we use a deep convolutional neural network on a publicly available dataset to extract descriptive features directly from the sequences, then fuse such feature with structural-and-physio-chemical driven features (such as amino-acid composition or AAIndex-based physicochemical properties). Dimentionality reduction is then performed on the resulting features and the output vectors are applied to train optimized gradient boosting machine (XGBoostt). We evaluate our method through three publicly available test sets, and show that our proposed DHS-Crystallize algorithm outperforms state-of-the-art methods, and achieves higher performance compared to using DCNN-deriven features, or structural-and-physio-chemical driven features alone.

## 1 Introduction

The current process of determining the protein’s structure mainly involves X-ray crystallography. However, such process is not only expensive with high attrition rate, but also has a very small success rate (ranging between 2 to 10 percent only, with more than 70% of the expenses relating to failed attempts) [13, 14]. To develop an alternative method, with less cost and attrition rate, number of previous studies have proposed machine learning based techniques [9, 1, 17, 16, 15, 12]. Such methods share the similar base-idea of analysing raw protein sequences and extracting physio-chemical, functional and sequence-based features from them. However, accurate estimation of protein crystallization propensity still remains a significant challenge.

Following the notable success of deep learning methods in many protein structure prediction [10, 15], and protein function prediction problems [7, 8, 10, 11]; recently a deep learning framework (Deep-Crystal) for sequence-based protein crystallization prediction was proposed [5].

Although the success of Deep-Crystal highlights the fact that such DCNN architect utilizes novel and complex non-linear-features from the provided raw protein sequences, it has been shown that a carefully handcrafted physio-chemical and sequence-derived features can result a much higher prediction [4].

Deep-Crystal uses deep learning framework to identify proteins which can produce diffraction-quality crystals, without manually engineering additional biochemical and structural features from sequence. Their result showed that using the raw protein sequences as an input of a Deep Convolutional Neural Network (DCNN) can outperform all other state-of-the-art sequence-based crystallization predictors.

Although the success of Deep-Crystal highlights the fact that such DCNN architect utilizes novel and complex non-linear-features from the provided raw protein sequences, it has been shown that a carefully handcrafted physio-chemical and sequence-derived features can result a much higher prediction [4].

In this paper, we introduce a hybrid method that takes advantage of both DCNN-driven features and physio-chemical-derived features (DHS-Crystallize) for accurate prediction of protein Crystallization. Our primary contribution includes:

- Fusing DCNN-driven with physio-chemical-derived features
- Performing Dimentionality reduction and training an optimized XGBoost modelby using resulted features
- Validate the method on three separate publicly available blind datasets, and proving that the proposed method can outperform the state-of-the-art methods

The rest of the paper is organized as following: Section.2 details the methodology behind DHS-Crystallize by describing the Datasets, feature extraction, and XGBoost classifier. We then evaluate our proposed DHS-Crystallize method and show the results in Section. 3. We also provide additional analysis on the performance of our physio-chemical-derived features and introduce our future work in 3. Finally, conclusion of our study is presented in Section.4.

## 2 Methodology

Predicting protein Crystallization is a binary classification task which requires an accurate labeling of the proteins with either crystallizable or non-crystallizable. In this section, we detail our proposed hybrid DHS-Crystallize (DHSC) method for predicting such property of proteins, based on their sequences.

### 2.1 Dataset

Following the same protocol as [5], our proposed DHSC method is trained and evaluated on publicly available datasets. For the training purpose, we have obtained the data from Wang et al. (2014) [17], that includes the total of 28731 proteins, with 5383 positive and 23348 negative samples.

Then, two blind test sets are obtained from SwissProt and Trembl databases (SP-final and TR-final), respectively. SP-final includes 148 crystallizable proteins and 89 non-crystallizable protein sequences. The second blind test set TR-final dataset, includes 374 crystallizable and 638 non-crystallizable.

Following the same protocol as [5], We further filter the sequences in training set and remove all the crystalizable protein sequences with similarity >15%, and all the non-crystalizable protein sequences with similarity >15%. That results in 2880 crystalizable protein sequences, and 8474 non-crystalizablle protein sequences in the final training set.

### 2.2 Feature Extraction

We start by extracting two type of features including DCNN-driven and Physio-Chemical-driven features, then fuse them to create the final descriptive feature for the protein sequences. We show in Section. 3 that the classifier built on the resulting features obtain better performance compared to using either of the two type of features alone.

#### 2.2.1 DCNN-driven features

To extract the DCNN-driven features, we employed the same DCNN architect described in [5] and pull the output of the fully connected layer to extract descriptive deep-network-based features (Fig. 1). To cover the effect of initialisation on the DCNN-driven-features, the network was trained 10 times, and the resulting features were concatenated, which made the final DCNN-based feature to be 2500 in length.

**Figure 1:**
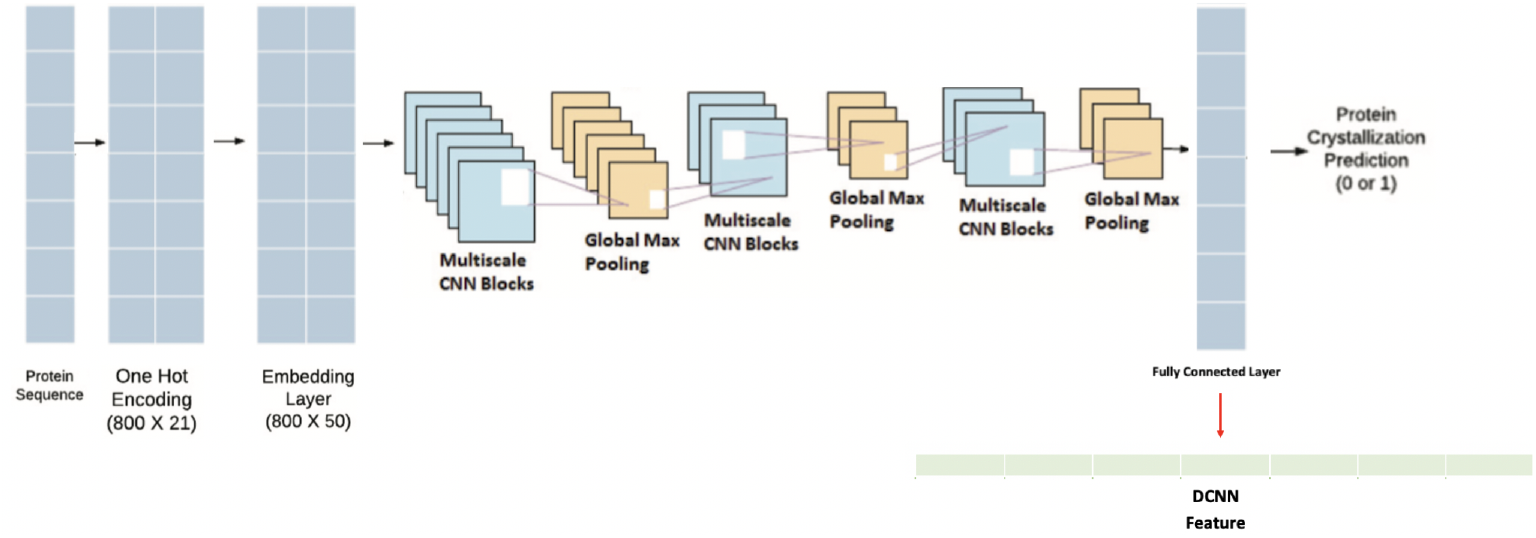
DCNN-Driven Feature

#### 2.2.2 Physio-Chemical-derived features

We have employed Protlearn software [3] to extract 21 Physio-Chemical-derived features from the provided protein sequences, as following:

- ***Length (LN)***: The number of amino acids that a protein is comprised of
- ***Amino Acid Composition (AAC)***: The frequency of amino acids for each sequence in the dataset
- ***AAIndex1-based physicochemical properties(AAindex1)***: A set of 20 numerical values representing various physicochemical and biological properties^1^ of amino acids. The indices are collected for each amino acid in the sequence.
- ***N-gram composition (NGram)***: We computes two set of features including both di- and tripeptide composition of amino acid sequences.
- ***Shannon entropy (entropy)***: Computes Shannon entropy for each sequence in the dataset as follows:

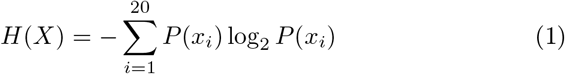

where i denotes the amino acid and *P* (*x*_*i*_) denotes the probability of a given amino acid in the sequence.
- ***Position-specific amino acids (posrich)***:Is a sparse vector where ones indicate the presence of the given amino acid(s) at the specified position(s), and zeros indicate their absence.
- ***Sequence motifs (motifs)***:binary vector indicating the presence of a specified amino acid sequence motif.
- ***Atomic and bond composition (ATC)***: calculates the sum of atomic and bond compositions for each amino acid sequence. Specifically, atomic features are comprised of five atoms: C, H, N, O, and S; and the bond features are comprised of the following: total bonds, single bonds, and double bonds.
- ***Binary profile pattern (binary)***: the binary profile pattern for each amino acid sequence in the dataset
- ***Composition of k-spaced amino acid pairs (CKAAP)***: the k-spaced amino acid pair composition of each sequence in the dataset.
- ***Conjoint triad descriptors (CTD)***: Initially developed to model protein-protein interactions, it groups Amino acids into 7 different classes based on their dipoles and side chain volumes (reflecting their electrostatic and hydrophobic interactions).
- ***Composition/Transition/Distribution - Composition (CTDC)***: The properties used here include: hydrophobicity^2^, normalized van der Waals volume, polarity, polarizability, charge, secondary structure, and solvent accessibility.
- ***Composition/Transition/Distribution - Transition (CTDT)***: Amino acids are categorized into 3 groups based on their physicochemical properties. The properties used here are same as those used for CTDC. This descriptor computes the frequency of transitions between groups as follows:

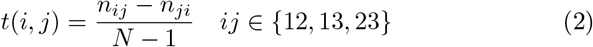

where *n*_*ij*_ and *n*_*ji*_ denote the frequency of transitions from group i to j and j to i, respectively, in the sequence, and N is the sequence length.
- ***Composition/Transition/Distribution - Distribution (CTDD)***: Contains grouped distribution of physicochemical properties. Since there are 13 physicochemical properties, 3 groups (see ctdc), and 5 distribution descriptors, the total dimensionality of this feature is 195.
- ***Normalized Moreau Broto autocorrelation*** ^3^ ***(moreau broto)***: Array containing Moreau-Broto autocorrelation descriptors.
- ***Moran’sI autocorrelation descriptors (Moran)***:Moran’sI descriptors are computed as follows:

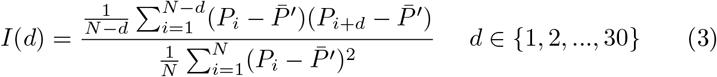

where *N*, *d*, *P*_*i*_, and *P*_*i*+*d*_ are the same as in the Moreau Broto autocorrelation; and 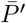 is the considered property P along the sequence, which is defined as follows:

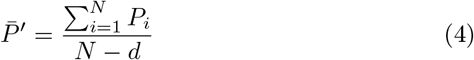
- ***Geary’s C autocorrelation descriptors (Geary)***: Geary’s C descriptors are computed as follows:

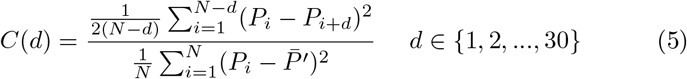

are *N*, *d*, *P*_*i*_, *P*_*i*+*d*_, and 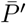 are the same as in the Moreau-Broto autocorrelation.
- ***Pseudo amino acid composition (PAAC)***: contains pseudo amino acid composition.
- ***Amphiphilic pseudo amino acid composition (APAAC)***: Contains amphiphilic pseudo amino acid composition
- ***Quasi-sequence-order (QSO*** : Contains QSO based on the Schneider-Wrede distance matrix.

### 2.3 Classification

For the classification purpose, we employ XGBoost algorithm [2], which is a saleable tree boosting method widely used by data scientists, as shown to provide state-of-the-art results on many problems. Tree boosting can be defined as a learning method designed to improve the classification of weaker classifiers through repeatedly adding new decision trees to the ensembles.

It has been shown by previous research works [6], that in cases when the dimentionality of the feature vector is very high, applying PCA before employing XGBoost can enhance both speed and performance of the XGBoost method. After applying the feature extraction method detailed in Sec. 2.2, each protein sequence is transformed into a descriptive vector of size 13078.

Therefore, we start by applying PCA to reduce the dimentionality of the feature vector to 300. Then, any classification technique can be employed; however, for the purpose of this study XGBoost was selected to classify the protein sequences into crystalizable or non-crystalizable. To find the best parameters for XGBoost, we have used 10% of the training data as validation set, and fine tuned our method. That resulted in the following setting to train the XGBoost algorithm: we used Learning rate of 0.01, maximum depth of 3, and the number of estimators of 500. To address the problem of imbalance dataset, we put more weight on the positive class which has less samples in the dataset, by setting the scale-pos-weight to 8.

The proposed pipeline is described in Fig. 2.

**Figure 2:**
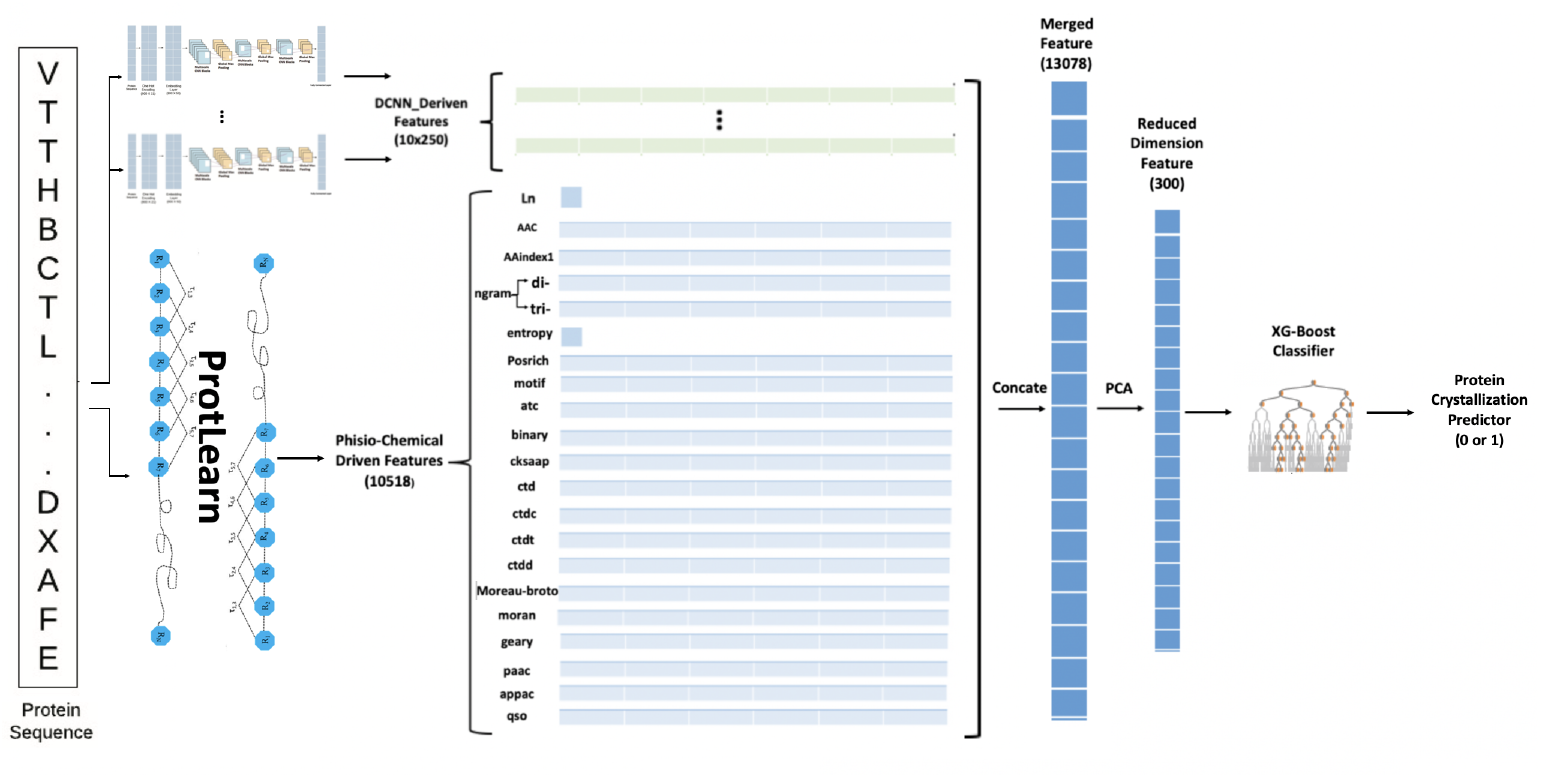
The pipeline of proposed DHS-Crystallize method

## 3 Result and Discussion

To evaluate the performance of the proposed DHS-Crystallize method, we have used three publicly available separate blind test sets, described in Sec. 2.1, namely: TR, SP, and Balanced. The results illustrated in bellow tables : Table 3, 2, and 1 shows that the proposed method outperforms state-of-the-art sequence-based methods, and leads to improved performance compared to using either Physio-Chemical-derived features or DCNN-driven features alone. Specifically, it leads to notable improvement on TR test set.

to evaluate the performance of Protlearn, we used the same setting as the one for DHSC (detailed in Section. 2.3). Thus, we started with reducing the features dimension to 300, which was following by employing XGBoost (with same setting as described in 2.3).

**Table 1:** Results obtained for protein crystallization prediction on the TR test data

| Methods | Accuracy | F1-Score | Precision | Recall |
| --- | --- | --- | --- | --- |
| CrysaliseII | 0.82 | 0.75 | 0.76 | 0.74 |
| PPCPred | 0.75 | 0.64 | 0.68 | 0.61 |
| CrystalP2 | 0.58 | 0.58 | 0.46 | 0.77 |
| DeepCrystal | 0.84 | 0.78 | 0.80 | 0.76 |
| Protlearn | 0.63 | 0.66 | 0.50 | **0.99** |
| DHSC | **0.89** | **0.86** | **0.81** | 0.92 |

**Table 2:** Results obtained for protein crystallization prediction on the SP test data

| Methods | Accuracy | F1-Score | Precision | Recall |
| --- | --- | --- | --- | --- |
| CrysaliseII | 0.75 | 0.78 | 0.86 | 0.72 |
| PPCPred | 0.67 | 0.67 | 0.86 | 0.55 |
| CrystalP2 | 0.66 | 0.73 | 0.71 | **0.76** |
| DeepCrystal | 0.76 | 0.79 | 0.88 | 0.72 |
| Protlearn | 0.75 | **0.82** | 0.95 | 0.73 |
| DHSC | **0.77** | 0.79 | **0.96** | 0.67 |

**Table 3:** Results obtained for protein crystallization prediction on the balanced test data

| Methods | Accuracy | F1-Score | Precision | Recall |
| --- | --- | --- | --- | --- |
| CrysaliseII | 0.8 | 0.8 | 0.83 | 0.77 |
| PPCPred | 0.67 | 0.62 | 0.74 | 0.53 |
| CrystalP2 | 0.58 | 0.63 | 0.57 | 0.7 |
| DeepCrystal | 0.83 | 0.82 | 0.85 | 0.79 |
| Protlearn | 0.71 | 0.77 | **0.98** | 0.64 |
| DHSC | **0.84** | **0.83** | 0.87 | **0.80** |

Tabel. 1 details the performance of the proposed DHSC on TR test sets, and shows that the proposed DHSC notably outperforms the state-of-the-art methods in protein crystallization on TR test set in accuracy, precision and F1-Score, but Protlearn achieves the highest Recall.

The performance of the method on SP is compared to other state-of-the-art algorithms and the results are reported in Table. 2. It demonstrates that the proposed DHSC outperforms the state-of-the-art methods in protein crystallization on SP test set in accuracy and precision, but Protlearn achieves the highest F1-Score and CrystallP2 achieves the highest Recall.

Table. 3 illustrates that the proposed DHSC outperforms the state-of-the-art methods in protein crystallization on balanced test set in accuracy and F-1 score, and Recall; but Protlearn achieves the highest precision.

The above tables also prove that merging the features leads to a better performance compared to only using Physio-Chemical-derived features or DCNN-driven features alone.

### 3.1 Additional Analysis and Future Work

As it can be observed from Table. 3, 2, and 1, protlearn performance is very imbalance when it comes to comparing Precision and Recall. We further investigated and found that a different setting for both PCA and XGboost can improve the Protlearn performance in all three TR, SP and Balanced dataset.

Specifically, following setting for PCA and XGBoost training algorithm led to highest protlearn performance : reducing the feature dimension to 15, using Learning rate of 0.001, maximum depth of 7, the number of estimators of 2000, and scale-pos-weight=3.5 to address the imbalance data set. The result of the analysis can be found in Figure. 3

**Figure 3:**
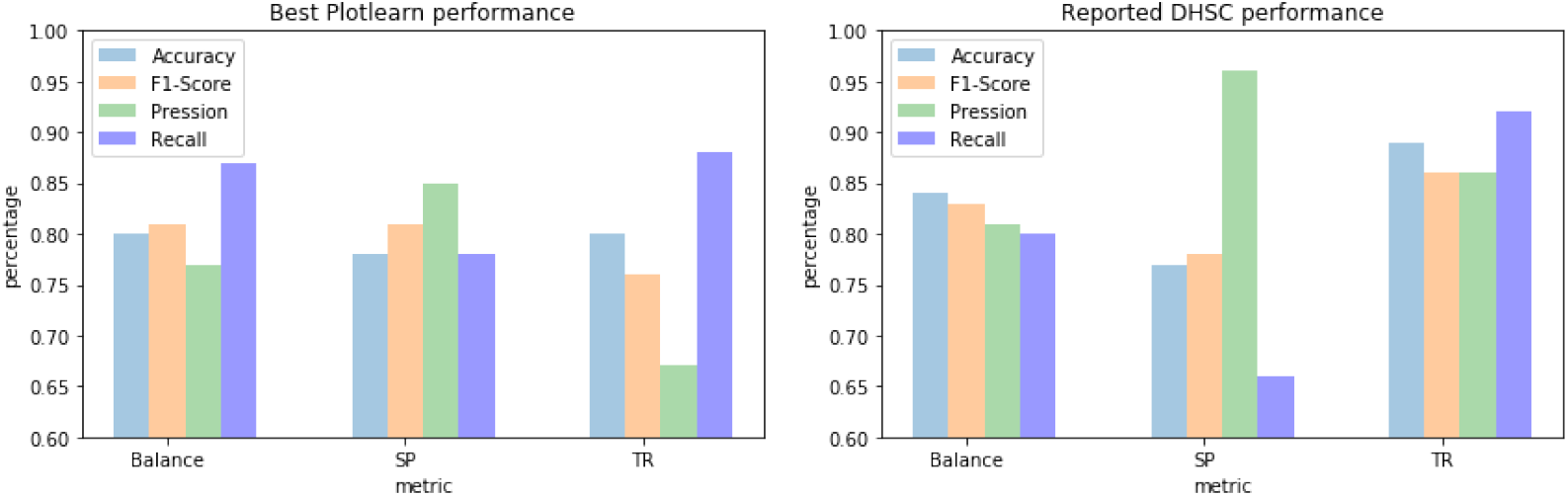
The best performance of protlearn

The result of using a lower dimension and optimized settings for the proposed DHS-Crystallize is yet to be investigated. However, it can be observed from Figure. 3, that the proposed DHS-Crystallize still have a better overall performance than fine-tuned Protlearn alone.

In addition, we find it non trivial to mention the most recent work on protein Crystallizability (BCrystall) based on only extracting Physio-Chemical-derived features which led to very high performance [4]. However, as the purpose of this study was to find if a hybrid model made of fusing DCNN-driven features and physio-Chemical-derived features would lead to advancement compared to using either of the features alone, we employed much simpler protlearn for computing physio-Chemical-derived features. As an extension to this work, we plan to investigate the affect of adding more physio-Chemical-derived features with respect to the latest investigation in BCrystall paper.

## 4 Conclusion

In this work, we investigated the effect of fusing DCNN-driven features with physio-Chemical-derived features on predicting crystalizability of the proteins. For this purpose, we have employed a deep convolutional neural network with state-of-the-art performance (DeepCrystall), and extracted the fully connected layer from the trained network to produce a descriptive feature for each protein sequence. To eliminate the effect of initialisation, we used 10 separately trained network and concatenated all the outputs. Then, we used protlearn software and extracted 21 physio-Chemical-derived features. We merged the resulting feature vectors and employed PCA to reduce the dimentionaloity and applied the resulting vectors to train XGBoost classifier. We have trained our model on a publicly available dataset and evaluated the performance of the proposed method on three blind test sets (also publicly available). The result indicates that the proposed method DHSC method outperforms all state-of-the-art algorithms and improves the performance comparing to when only one set of features are used.

## Acknowledgments

This work was supported by an Investigator Grant from the National Health and Medical Research Council (NHMRC) of Australia (GNT1174405) and the Victorian Government’s OIS Program.

## Footnotes

1 ver.9.2 (AAindex1 release Feb, 2017)

2 For hydrophobicity, seven different groupings were used (based on different studies), which can all be found in AAIndex1.

3 based on AAIndex1

## References

[1] Phasit Charoenkwan et al. “SCMCRYS: predicting protein crystallization using an ensemble scoring card method with estimating propensity scores of P-collocated amino acid pairs”. In: PloS one 8.9 (2013), e72368.

[2] Tianqi Chen and Carlos Guestrin. “Xgboost: A scalable tree boosting system”. In: Proceedings of the 22nd acm sigkdd international conference on knowledge discovery and data mining. 2016, pp. 785–794.

[3] Thomas Dorfer. Protlearn. Version 1. Oct. 2, 2020. url: https://github.com/tadorfer/protlearn.

[4] Abdurrahman Elbasir et al. “BCrystal: an interpretable sequence-based protein crystallization predictor”. In: Bioinformatics 36.5 (2020), pp. 1429–1438.

[5] Abdurrahman Elbasir et al. “DeepCrystal: a deep learning framework for sequence-based protein crystallization prediction”. In: Bioinformatics 35.13 (2019), pp. 2216–2225.

[6] Nilam Fitriah et al. “EEG channels reduction using PCA to increase XGBoost’s accuracy for stroke detection”. In: AIP Conference Proceedings. Vol. 1862. 1. AIP Publishing LLC. 2017, p. 030128.

[7] Sameer Khurana et al. “DeepSol: a deep learning framework for sequence-based protein solubility prediction”. In: Bioinformatics 34.15 (2018), pp. 2605–2613.

[8] Maxat Kulmanov, Mohammed Asif Khan, and Robert Hoehndorf. “DeepGO: predicting protein functions from sequence and interactions using a deep ontology-aware classifier”. In: Bioinformatics 34.4 (2018), pp. 660–668.

[9] Lukasz Kurgan, Marcin J Mizianty, et al. “Sequence-based protein crystallization propensity prediction for structural genomics: review and comparative analysis”. In: Natural Science 1.02 (2009), p. 93.

[10] Zhen Li and Yizhou Yu. “Protein secondary structure prediction using cascaded convolutional and recurrent neural networks”. In: arXiv preprint arXiv:1604.07176 (2016).

[11] Raghvendra Mall et al. “An unsupervised disease module identification technique in biological networks using novel quality metric based on connectivity, conductance and modularity”. In: F1000Research 7.378 (2018), p. 378.

[12] Fanchi Meng, Chen Wang, and Lukasz Kurgan. “fDETECT webserver: fast predictor of propensity for protein production, purification, and crystallization”. In: BMC bioinformatics 18.1 (2017), p. 580.

[13] Robert Service. Structural genomics, round 2. 2005.

[14] Thomas C Terwilliger, David Stuart, and Shigeyuki Yokoyama. “Lessons from structural genomics”. In: Annual review of biophysics 38 (2009), pp. 371–383.

[15] Huilin Wang et al. “Critical evaluation of bioinformatics tools for the prediction of protein crystallization propensity”. In: Briefings in bioinformatics 19.5 (2018), pp. 838–852.

[16] Huilin Wang et al. “Crysalis: an integrated server for computational analysis and design of protein crystallization”. In: Scientific reports 6 (2016), p. 21383.

[17] Huilin Wang et al. “PredPPCrys: accurate prediction of sequence cloning, protein production, purification and crystallization propensity from protein sequences using multi-step heterogeneous feature fusion and selection”. In: PloS one 9.8 (2014), e105902.

